# Spectral variation and pigmentary basis of ornamental and mimetic wing colour patches of swallowtail butterflies

**DOI:** 10.1101/2024.01.28.577613

**Authors:** Bhavya Dharmaraaj, Radhika Venkatesan, Krushnamegh Kunte

**Affiliations:** National Centre for Biological Sciences, Tata Institute of Fundamental Research, GKVK Campus, Bellary Road, Bengaluru 560065, India; Indian Institute of Science Education and Research Kolkata, Mohanpur, Nadia, West Bengal 741 246, India

**Keywords:** Animal signals, butterfly wing patterns, colour properties, sexual dimorphism

## Abstract

Colours and colour patterns are extraordinarily diverse traits that are often used as visual signals. To test ecological and evolutionary drivers of these visual signals, a clear understanding of their nature and variation is necessary. Here we characterise variation in wing colouration of Asian mormon swallowtail butterflies (*Papilio*, subgenus *Menelaides*). These species exhibit two kinds of colour patches on largely black wings: creamy white/yellow/green patches that are presumably used as sexual ornaments, and pure white patches that are presumably used as mimetic signals. Using reflectance spectrophotometry we quantified spectral properties of black wing background and colour patches between sexes, wing surfaces and mimicry status. We discovered that brightness and saturation of the black background were less variable across sexes, wing surfaces and mimetic/non-mimetic status. However, colour contrast and saturation were higher on dorsal surfaces than on ventral surfaces, and colour contrast between the black background and colour patches was higher in males than in females. Colour patches in non-mimetic butterflies were brighter and more saturated across the whole colour spectrum compared to mimetic butterflies. These patterns of colour variations in relation to their putative functions suggest that: (a) colour patches on dorsal and ventral wing surfaces evolve independently to accommodate differential strengths of natural and sexual selection, (b) sexual ornaments are brighter in non-mimetic males but they also occur in all non-mimetic females, indicating mutual sexual selection on these ornaments, but which is stronger in males, and (c) mimetic male and female butterflies display less sexual contrast in colour patches, indicating relatively similar strength of mimetic selection. Thus, our study characterises colour variation in an extraordinary signal radiation on the wings of swallowtail butterflies, a model clade in ecology, evolution and genetics. Finally, using liquid chromatography-mass spectrometry (LCMS) we identified the pigment papiliochrome-II to be the chemical basis of the presumed sexual ornaments in mormon swallowtails.

## Introduction

Colour patterns of animals exhibit immense variation between sexes and across species. The adaptive functions of colour patterns include thermoregulation (Stuart-Fox et al. 2017), predator avoidance (Ruxton et al. 2004), protection from radiation (Salih et al. 2000), and courtship (Pryke and Griffith 2007). These functions indicate that colour patterns are subject to both natural and sexual selection either acting antagonistically or synergistically to shape colour variation. This can result in colour patterns being used under different functional contexts (Estrada and Jiggins 2008). For example, poison arrow frog colour patterns are shaped by both natural and sexual selection as they are used in antipredator defences in males and females as well as courtship by males (Maan and Cummings 2008; Noonan and Comeault 2009). Similarly, butterfly wings colours are also multifunctional, often resulting from a balance between courtship, defensive colouration and thermoregulation (Ellers and Boggs 2003; Estrada and Jiggins 2008). These selective regimes can affect colouration differently in males and females and can bring about sexually dichromatic phenotypes (Stuart-Fox and Ord 2004; Heinsohn et al. 2005; Dale et al. 2015). The evolution of sexual dimorphism in colouration has been explored primarily through two alternative models i) Darwin’s model where sexual selection acts on males through female choice, resulting in male traits diverging from ancestral phenotypes and males evolving elaborate sexual ornaments or colouration (Darwin 1871); and ii) Wallace’s model where natural selection acts on females, resulting in female traits diverging to evolve protective or cryptic colouration whereas males retain the ancestral phenotype (Wallace 1897). Studies have reported evidence for both models affecting colour evolution across taxa (Badyaev and Hill 2003; Oliver and Monteiro 2011; Shultz and Burns 2017; Cooney et al. 2019). Owing to their immense colour diversity, butterflies are one of the major study systems for exploring the evolution of sexual dimorphism in visual cues, in which evidence for both models has accumulated (Kunte 2008; van der Bijl et al. 2020).

In butterflies, differential selective pressures can lead to variation in wing colours between males and females and along the dorsoventral axis within individuals. This variation can include complete changes in wing colour patterns (Kunte 2008; Prakash and Monteiro 2018) or more subtle continuous variation in colour intensity or quality; e.g., where one sex can appear darker (Tuomaala et al. 2012). The widespread variation of colour patterns between sexes and across wing surfaces has generated multiple ecological and evolutionary hypotheses to explain functions and evolutionary trajectories of butterfly wing colouration. For instance, variation in the dorsal UV reflective patches in male *Colias eurytheme* and *Eurema hecabe* butterflies were found to influence female mate preference indicative of their function as sexual ornaments (Kemp and Rutowski 2011). Butterflies of another coliadine species, *Colias philodice eriphyle*, do not reflect UV. However, both sexes of *Colias philodice eriphyle* and *Colias eurytheme* vary in ventral melanisation levels, further affecting processes such as thermoregulation, egg maturation rates, and mate-choice, which is a good example of how wing colouration may evolve to accommodate varied aspects of natural and sexual selection (Ellers and Boggs 2003, 2004). Along with the use of wing colour patterns as signals in intraspecific communication (courtship, species recognition), colour patterns are also used in interspecific communication, e.g., in protective colouration under aposematism and mimicry (Estrada and Jiggins 2008; Finkbeiner et al. 2014). Signals can be partitioned along the dorsoventral axis to alleviate antagonistic selection pressures—the dorsal surface often bearing sexual ornaments or flashing colour patches/eyespots that may startle predators, while the ventral surface often being used in protective colouration such as for crypsis (Vallin et al. 2005; Oliver et al. 2009; Merilaita et al. 2011; Su et al. 2015).

Under the paradigm of Batesian mimicry, which is an adaptation in which a palatable species (the mimic) gains a level of protection against predators by resembling an aposematic species (the model) (Ruxton et al. 2004), butterfly wing colour patterns have diverged considerably where wing colours evolve at faster rates than other traits (Basu et al. 2023). Batesian mimicry varies in complexity from monomorphic to dimorphic and polymorphic mimicry, and can also be limited to females while males retain the non-mimetic phenotype (Kunte 2009b,a). *Papilio* swallowtails are a diverse group of butterflies with widespread mimicry and wing pattern variation. While several studies have addressed the genetics of mimicry in a few *Papilio* species (Koch et al. 2000; Kunte et al. 2014; Timmermans et al. 2014; Iijima et al. 2018), the chemical basis of colour patterns (i.e., papiliochrome II) in others (Umebachi 1985; Nishikawa et al. 2013; Yoda et al. 2021), and the extent of mimetic resemblance in a few species pairs (Yoda et al. 2021), studies exploring the natural variation between different colour patches itself is lacking. This is of relevance as subtle variations in colours have been implicated in various phenomena such as species recognition (Fordyce et al. 2002), mate choice and mating success (Davis et al. 2007) and mimicry (Rossato et al. 2018). *Menelaides* – a species-rich subgenus of *Papilio* – contains the largest number of Batesian mimics within the genus (Kizhakke and Kunte 2022; Condamine et al. 2023). Several *Menelaides* species exhibit female-limited or female-limited polymorphic mimicry, resulting in a spectacular diversity of wing colours between and within sexes (Kunte 2009a; Kizhakke and Kunte 2022). Colour patterns of these butterflies predominantly contain melanised (black or dark brown) wings as a background colour on which white, yellow, green, blue or red colour patches are placed on both dorsal and ventral surfaces. Along with mimetic dimorphism and polymorphism, there exists intraspecific variation in the white patches where some possess creamy white/yellow/green patches whereas others possess pure white patches. Moreover, we observed that these patches also appear to vary in intensity of colouration, along the dorsoventral axis and between males and females.

Using this information on wing colour variation, we first quantified the spectral differences between male and female *Menelaides*, their dorsal and ventral surfaces, and individuals with different mimetic status. We use reflectance spectrophotometry to parse the variation in wing colour patches and background black colour. This method facilitates quantification of colours and extract their spectral parameters such as brightness, hue, and saturation (Endler 1990). We characterized the dark wing background and bright colour patches to quantify variation that contributes to sexual dimorphism and dorsoventrally mismatched colouration, and to address the following questions: (1) Are colour patches of male butterflies brighter and more saturated than patches of female?, (2) Do both sexes show similar contrast between black background and colour patches (potential visual signals), i.e., between background and signal patches that affects conspicuousness and efficacy of colour signals?, (3) Are colour patches on dorsal wing surfaces that are often used in courtship, brighter and more saturated than those on ventral surfaces?, (4) How is colour variation structured when the colour patches are used in mimicry rather than in courtship? Finally, we performed chemical analysis using liquid chromatography-mass spectrometry (LCMS) to characterise the mechanistic basis of colouration in these colour patches. This is a very broad characterization of wing colour variation in this super-diverse subgenus. We use this characterization of sex- and wing surface-specific colour variation to generate specific hypotheses in the context of butterfly behaviour and communication. Our work provides a clear path to more hypothesis-driven studies of evolution of colour patterns in this model clades that includes iconic polymorphic species such as *Papilio polytes* and *P. memnon*.

## Materials and Methods

### (a) Reflectance Measurements

To characterize colour, we measured reflectance spectra of museum specimens of *Menelaides* butterflies using an Ocean Optics Jaz spectrometer equipped with a pulsed Xenon lamp (PX-1 lamp). We measured reflectance of forewing and hindwing black and white/yellow/green patches on dorsal and ventral surfaces. For each dataset, we measured only specimens in good condition; i.e., those with no perceptible fading of colour and with little wing wear and tear, and wherever possible, recently collected specimens, to generate a high-quality dataset with reliable colour measurements. We chose specimens carefully, ensuring that they were pinned properly; i.e., wings lay flat and invariably in a plane with the body, so that the photospectrometer readings were taken with the desired angle of light and collecting fibre. To collect spectral measurements, we used two optic fibres fitted with collimating lenses. We placed the illuminating probe at 90° and the collecting fibre at 45° to the wing surface. We measured reflectance with respect to a Spectralon white standard which reflects 96% of incident light. We set the diameter of the illuminating spot to be 1 mm, which was much smaller than sizes of colour patches. We enclosed the setup in a cardboard box covered with black felt-cloth to exclude ambient light. We took 1–3 readings of the colour patches on both wing surfaces and averaged them, using the R package Pavo (Maia et al. 2019), and used the averaged spectra for all downstream analysis (see Tables S1 and S2).

We used two datasets separately to compare colour variation: 1) A dataset (referred to as ‘Dataset 1’ in this work) of measurement of reflectance spectra from one male and one female of *Menelaides* species. In polymorphic species, we measured one individual each of the different morphs. This dataset included 47 species and additional six morphs (see Table S1 for species/morph details). It was not possible due to time constraints and for practical and economic reasons to measure multiple males and females of each *Menelaides* species. 2) A dataset (referred to as ‘Dataset 2’ in this work) containing 3–4 species and 4–5 individuals of each sex, of mimetic and non-mimetic categories depending on the availability of museum specimens (females of most *Papilio* are under-represented in museum collections, just as they are less frequently encountered in nature compared to males). The carefully chosen specimens for Dataset 1 ensured that the measurements on single individuals of each sex would be representative of the species, and this dataset offered a broader window into the properties of colour patches at the entire clade level. More intensive sampling of select species/morphs in the non-mimetic and mimetic categories in Dataset 2 ensured that a measure of inter-individual variation was considered in the more focused analysis. Thus, the two datasets were compositionally distinct and complementary in addressing the questions raised in this work.

### (b) Colour Descriptors

From the averaged spectra for black and white colours, we extracted colour variables for brightness and saturation. We used the total brightness, which is the area under the curve, (termed B1 from (Maia et al. 2019)) of the patch to characterize brightness. As black and white spectra do not have a single peak of reflectance, we used the difference between maximum and minimum reflectance with respect to the average brightness as saturation (termed S8 (Maia et al. 2019)). Further, we calculated brightness and chroma in the UV region as total reflectance from 300 nm–400 nm (UVB3), and reflectance in the UV region divided by the total reflectance (S1U), respectively. We also calculated a contrast ratio between the black/brown wing background and white/yellow/green patches, as brightness of the patches divided by brightness of the black/brown background. Thus, a higher ratio indicates more contrast between the patches, which may be treated as conspicuousness of the colour patches.

### (c) Data Curation and Statistical Analysis

We parsed the data into four categories based on sex and wing surface, as (a) male-dorsal, (b) male-ventral, (c) female-dorsal, and (d) female-ventral. We performed pairwise comparisons in colour parameters between: (1) the same wing surfaces of males and females; and (2) dorsal and ventral surfaces of individuals of the same sex (See Table S1 for details of sample sizes and comparisons). We performed statistical analysis of Dataset 1 and Dataset 2 separately, and their results are also represented separately in tables and figures below.

Next, using only the Dataset 1, we divided males and females into mimetic and non-mimetic categories. We then calculated: (a) pairwise differences in dorsal versus ventral wing surfaces in the white patches of mimics and non-mimics (Fig. 3), (b) differences of black/brown versus white/yellow/green patches on the same wing surface between mimetic and non-mimetic males and females (Fig. 4, S4), and (c) differences in spectral parameters for black/brown and white/yellow/green patches on the same wing surface in mimics versus non-mimics in both sexes (Fig. 4, S4). We carried out statistical analyses in R v.4.0.4. We used Shapiro-wilk tests to assess normality. Depending on whether the data were normally or non-normally distributed, we compared groups using paired sign tests from the R package *rstatix* (Kassambara 2023), *t*-tests, or Mann-Whitney tests, as appropriate.

Since black wing patches reflect much lesser amount of light than white wing patches, we scaled all the spectral parameters between 0 and 1 using the ‘rescale’ function from the *scales* package (Wickham 2022). We did this normalization independently for black and white patches so that the normalized values could be compared with each other for the differential ranges across colours, wing surfaces, sexes and mimicry status.

### (d) Chemical Characterization of Creamy White/Yellow/Green Patches and Pure White Patches

To chemically characterize wing colouration in *Papilio*, we performed solvent extraction of pigments from the hindwing white patches of two species: 1) *Papilio polytes* (a species that exhibits female-limited polymorphic Batesian mimicry, i.e., its males and one female form are non-mimetic, and two other female forms are mimetic), and 2) *Papilio demoleus* (a non-mimetic, sexually monomorphic species) (Fig. 5A). We separately cut out hindwing patches of three males each of *P. polytes* and *P. demoleus* (non-mimetic butterflies) containing the creamy white/yellow colouration and three females of *P. polytes f. polytes* (mimetic butterflies) for pure white colouration. We used 1 ml of 100% HPLC-grade methanol as a solvent for extraction. We sonicated and then filtered the solution using a membrane filter of 0.12 µm. We used the filtrates of the three samples for LCMS analysis.

For LCMS analysis, we used a Dionex Ultimate3000 UHPLC coupled with Thermo Fisher Q Exactive for separation and identification of compounds. We used a C18 column (Phenomenex Luna) with particle size 5 µm and dimensions of 150 mm x 4.6 mm. The elution gradient (time in minutes [% solvent B]) was as follows: 0–2[0.2%], 2–20[0.2–20%], 20–35[20–60%], 35–40[60–100%], 40– 45[100%], 45–45.1[95–0.2%], 45.1–55[0.2%]. Solvent A was 10 mM Ammonium acetate in water (0.1% formic acid) and solvent B was 0.1% formic acid in Acetonitrile. We maintained the flow rate at 0.4 ml/min. For the mass analysis we first scanned for the mass of papiliochrome-II in all the samples. From these primary mass spectra, we also identified a major product ion, and we carried out a scan monitoring for both papiliochrome-II and the product ion across the samples. We normalised intensities across samples to the sample with the highest intensity of papiliochrome-II to get the relative abundance.

## Results

### Colour Patches in Males Were Brighter, More Saturated, and With Higher Contrast, Than Those in Females

We compared the black/brown background and colour patches of males versus females to assess differences in brightness and saturation (Fig. 1A). In both Datasets 1 and 2, black/brown background colour was darker (less bright) in males than in females on both dorsal and ventral wing surfaces (statistical details in Table S3). Males had brighter colour patches irrespective of wing surface (Fig. 1B, D). To quantify conspicuousness, we calculated the contrast ratio between the black/brown background and colour patches. Contrast between patches was higher in males than in females (**Dataset 1:** Welch two-sample t-test for the dorsal surface: *t*=-2.898, *df*=30.029, *p*=0.007, Welch two sample t-test for the ventral surface: *t*=-2.1797, *df*=33.13, *p*=0.04, Fig. 1C. **Dataset 2:** Wilcoxon rank sum test for the dorsal surface: *W*= 62, *p*=0.0003, Wilcoxon rank sum test for the ventral surface: *W*= 108, *p*=0.03, Fig. 1E), indicating that males had more conspicuous colour patches on a darker background compared with females. Black/brown background and colour patches on ventral wing surfaces were more saturated in males than in females in Dataset 1, but not in Dataset 2 (Fig. S1A, B). Therefore, colour saturation did not vary between the sexes on dorsal wing surfaces but males had more saturated colour patches on ventral wing surfaces compared with females (Table S3).

**Figure 1:**
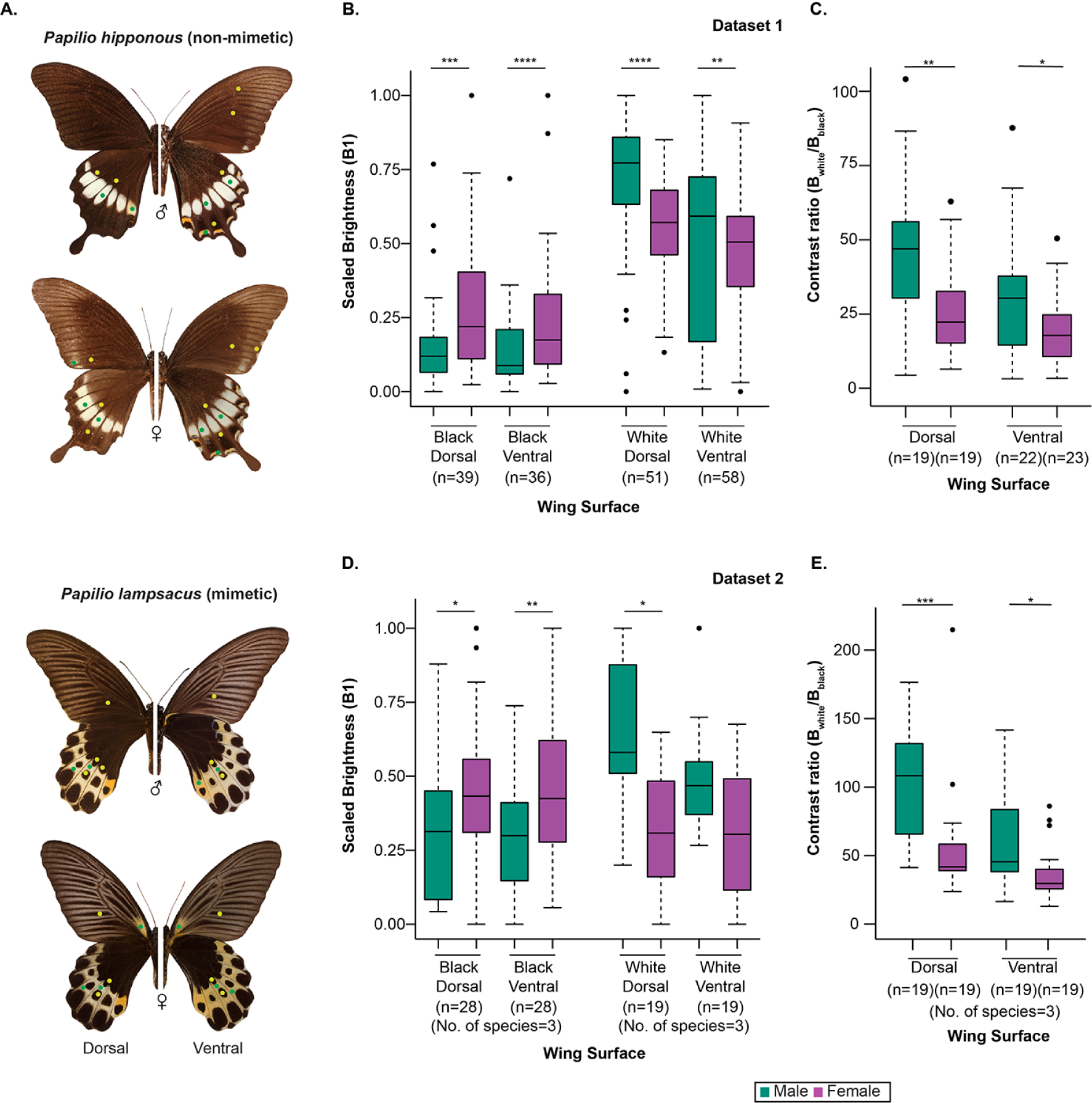
Comparison of dark wing background and colour patches on the wing surfaces of swallowtail butterflies (*Papilio*). **A:** Representative images of males and females of a non-mimetic and mimetic species: spots indicate wing areas where spectral readings for black/brown (yellow spots) and white patches (green spots), were taken. **B–C: Dataset 1**, comprising one individual each of male and female of all *Papilio* (*Menelaides*) species**, D–E: Dataset 2:** multiple individuals each of male and female of a selection of *Papilio* (*Menelaides*) species. **B and D:** Comparison of paired values of black/brown and white patches across sexes on the same wing surface. Brightness values are normalized from 0 to 1 based on min-max readings. **C and E:** Comparison of contrast (raw values of brightness of white/brightness of black) between males and females on the same wing surface. Black dots represent outliers. Statistical significance of pairwise sign tests: *: p<0.05, **: p<0.01, ***: p<0.001, ****: p<0.0001.

### Dorsal Wing Surfaces Had More Saturated Colour Patches, Ventral Wing Surfaces Had Brighter and More UV Reflective Colour Patches

We expected based on predictions of sexual and natural selection theory for black/brown wing backgrounds to be more saturated and colour patches to be brighter on dorsal wing surfaces than on ventral surfaces. This is because dorsal surfaces are typically used in courtship whereas ventral surfaces are often believed not to play a major role in courtship in *Papilio* and many other butterflies, which typically have more cryptic or otherwise less conspicuous ventral sides (Lederhouse and Scriber 1996; Kemp and Macedonia 2006; Oliver et al. 2009). Contrary to these expectations, colour patches showed similar levels of brightness and saturation on both wing surfaces in both datasets. In males, black/brown backgrounds were darker on dorsal surfaces but the colour patches did not differ across wing surfaces, whereas in females black/brown background and colour patches were brighter on ventral surfaces (Table S4, Fig. 2A, C). In colour patches of males, dorsal surfaces had higher contrast than ventral surfaces (**Dataset 1:** Wilcoxon rank sum test for males, *W*= 1369, *p*=0.0004, Fig. 2B, **Dataset 2:** Wilcoxon rank sum test for males, *W*= 461, *p*=0.003, Fig. 2D). Females also had higher contrast on dorsal surfaces, though significantly different than the ventral surface only in Dataset 2 (**Dataset 2:** Wilcoxon rank sum test for females, *W*= 416, *p*=0.007, Fig. 2D). Saturation of the black/brown background was higher on the dorsal surface than ventral surface in females in both datasets, whereas saturation did not differ in males (Table S4, Fig. S2). Colour patches were more saturated on dorsal surfaces than on ventral surfaces in both sexes (Table S4, Fig. S2). Thus, dorsal surfaces had more saturated colour patches whereas ventral surfaces had brighter patches (Table S4).

**Figure 2:**
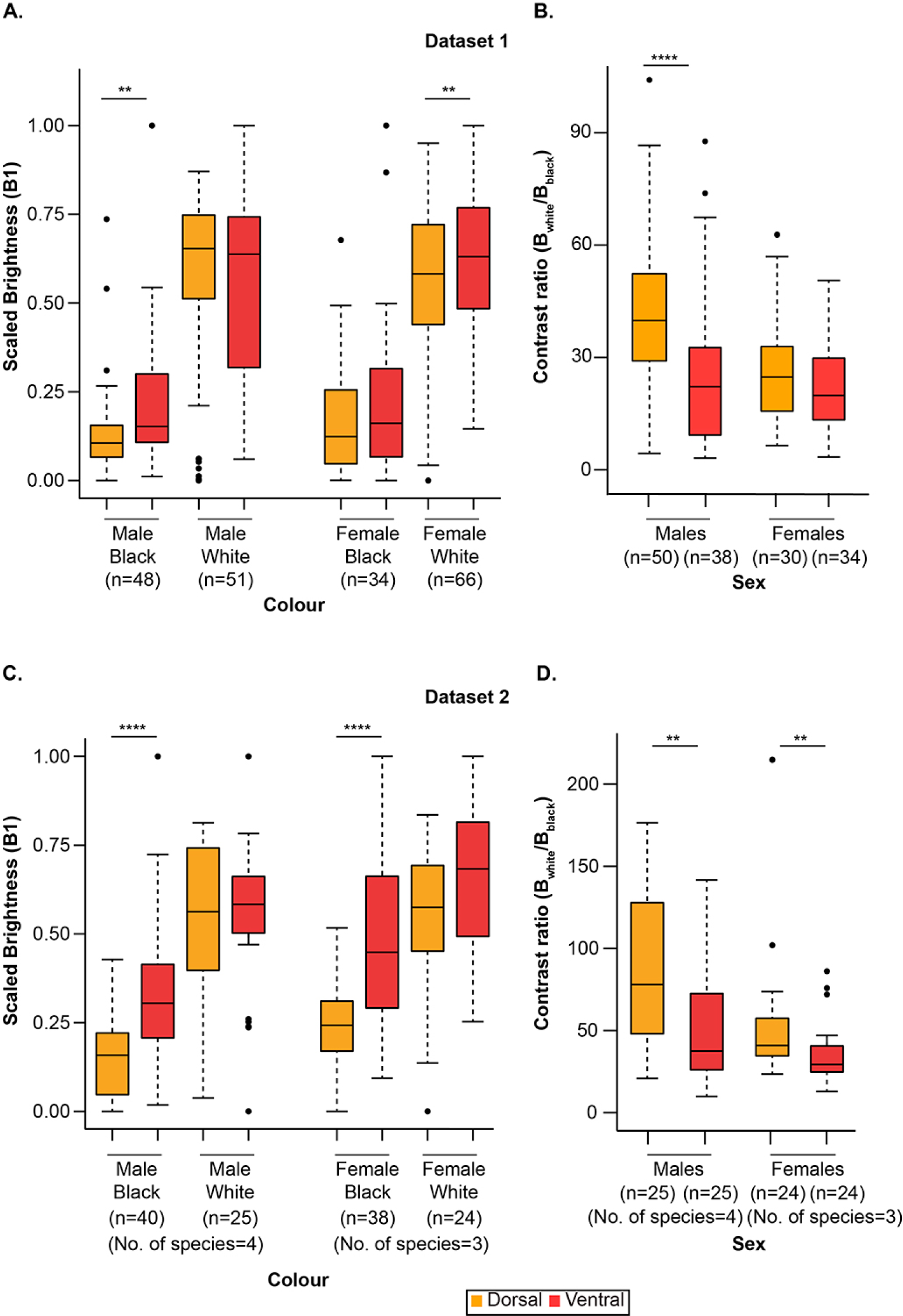
Comparisons of black/brown and white colours within sex across wing surfaces. **A–B:** Dataset 1; **C–D:** Dataset 2. **A and C:** Comparison of paired values of dorsal and ventral black/brown and white patches. Brightness values are normalized from 0 to 1 based on min-max readings. **B and D:** Comparison of contrast (raw values of brightness of white/brightness of black) along dorsoventral axis within each sex. Black dots represent outliers. Brightness values are normalized from 0 to 1 based on min-max readings. Black dots represent outliers. Statistical significance of pairwise sign tests: *: p<0.05, **: p<0.01, ***: p<0.001, ****: p<0.0001.

Most mimetic *Menelaides* exhibit female-limited mimicry, and only a few species exhibit sexually monomorphic mimicry. Creamy white/yellow/green patches tended to be present in non-mimetic species and colour forms, whereas bright, pure white patches were usually present in mimics. Reflectance spectra of colour patches differed between mimics versus non-mimics, with mimics showing a prominent peak in the UV range (300–400 nm; see Fig. S3 for reflectance spectra) that was absent in most non-mimics. To characterise overall colour variation, we analysed brightness and saturation for the whole light spectrum (300–700 nm) and then separately just in the UV range.

In the analysis of whole light spectrum, brightness was comparable on dorsal and ventral wing surfaces, but saturation was higher on dorsal compared with ventral wing surfaces (Table S5, Fig. 3A– B, and for details of pairwise comparisons see Table S9B, green boxes). On the other hand, analysis of only the UV range showed that UV brightness (UVB3) and UV saturation (S1U) were higher on ventral surfaces (Table S5, Fig, 3C–D, Table S9C). This indicated that ventral wing surfaces reflected more UV in mimetic butterflies even when the total brightness in the whole colour range was not affected (Fig. 3A). In contrast, colour patches of non-mimics were more saturated on dorsal surfaces but brighter and more UV reflective on ventral surfaces, although less UV reflective (Fig. 3A and 3C). UV reflectance affects the visual appearance of colour patches, and the stark difference in UV reflectance across wing surfaces in mimics and non-mimics suggests that signal function and efficacy may vary depending on wing surfaces on which colour patches are present (Table S5, Fig. S3). It is likely that UV reflectance itself may be subject to selection independently of reflectance across the whole spectrum.

**Figure 3:**
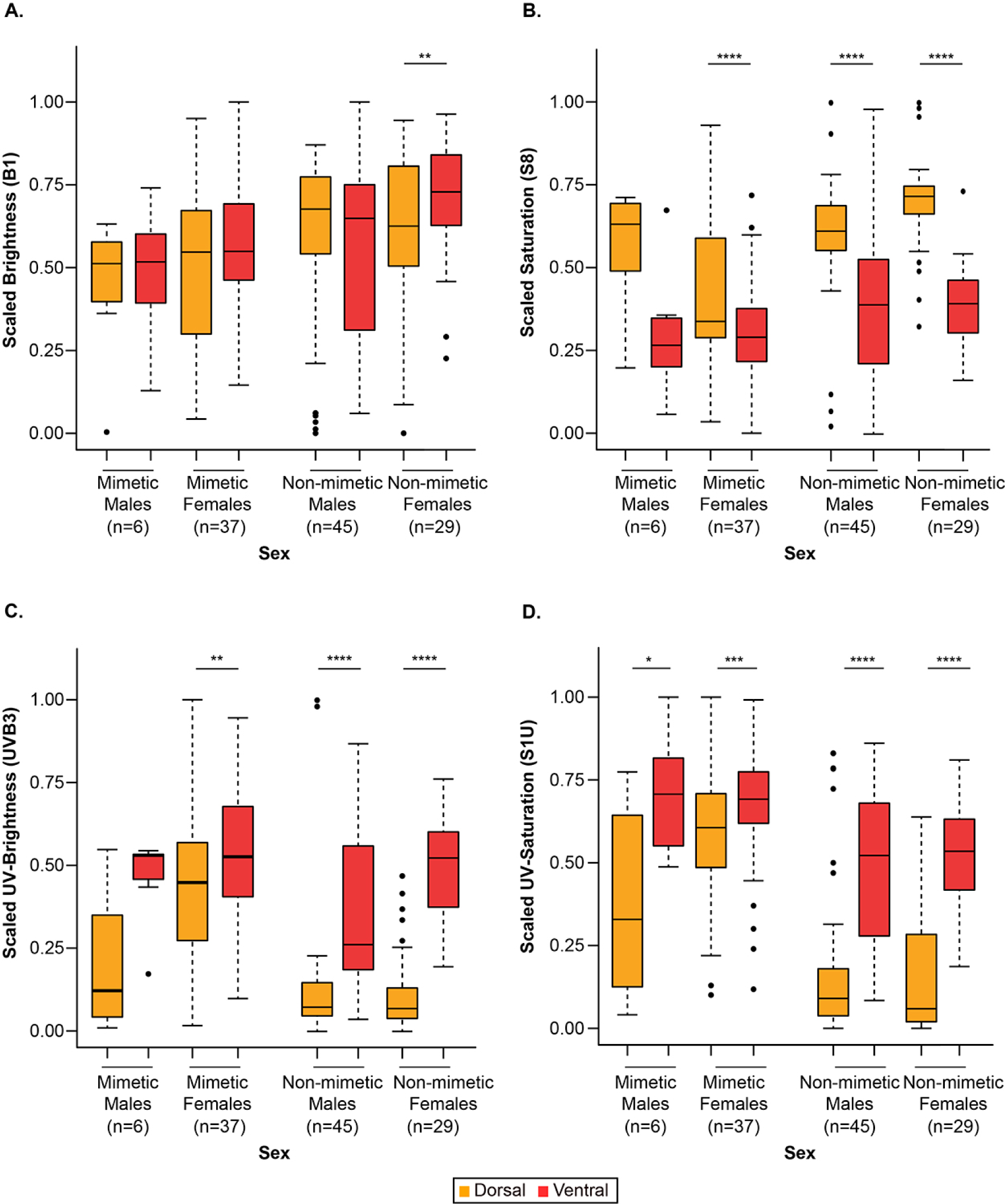
Comparison of colour patches on different wing surfaces of mimics and non-mimics. **A:** brightness, and **B:** saturation, of colour patches within mimics and non-mimics on two wing surfaces across the entire spectral range (300 nm–700 nm). **C:** UV-brightness, and **D:** UV-chroma of colour wing colour patches in mimics and non-mimics in the UV range (300 nm–400 nm). Brightness values are normalized from 0 to 1 based on min-max readings. Black dots represent outliers. Statistical significance of pairwise sign tests: *: p<0.05, **: p<0.01, ***: p<0.001, ****: p<0.0001.

### Black/Brown Background Did Not Vary in Saturation, but it was less dark in Non-Mimetic Females

There was no significant difference in the saturation of black/brown wing background colouration between mimetic and non-mimetic males or females, except that non-mimetic females had less dark background on the ventral surface (Table S6A, Fig. S4A–B, and for details of pairwise comparisons see Table S9A, green boxes).

We then compared black/brown background between mimetic males and females, and non-mimetic males and females. The non-mimetic females had less dark background than their non-mimetic male counterparts, while saturation showed no differences between the sexes (Fig S4A–B, orange lines and stars, Table S7A, and for details of pairwise comparisons see Table S9A, yellow boxes). This indicated that non-mimetic females had brighter background colouration, predominantly on ventral wing surfaces.

### Non-Mimetic Females Had More Saturated as well as Brighter Colour Patches Than Those of Mimetic Females

Ventral colour patches of non-mimetic females were brighter than those of mimetic females (Table S6B, Fig. 4A, black lines and stars, and for details of pairwise comparisons see Table S9B, blue boxes). On the other hand, non-mimics tended to have brighter colour patches in both sexes than those of mimics, but not significantly so (Fig. 4A). Non-mimetic females also had more saturated colour patches than mimetic females (Table S6B, black lines and stars in Fig. 4B, blue boxes in Table S9B). Saturation and brightness of non-mimetic and mimetic males did not differ (Fig. 4A, B). Further, as expected, mimics had higher UV reflectance and UV saturation than non-mimics (Table S6B, black lines and stars in Fig. 4C–D, and blue boxes in Table S9C).

**Figure 4:**
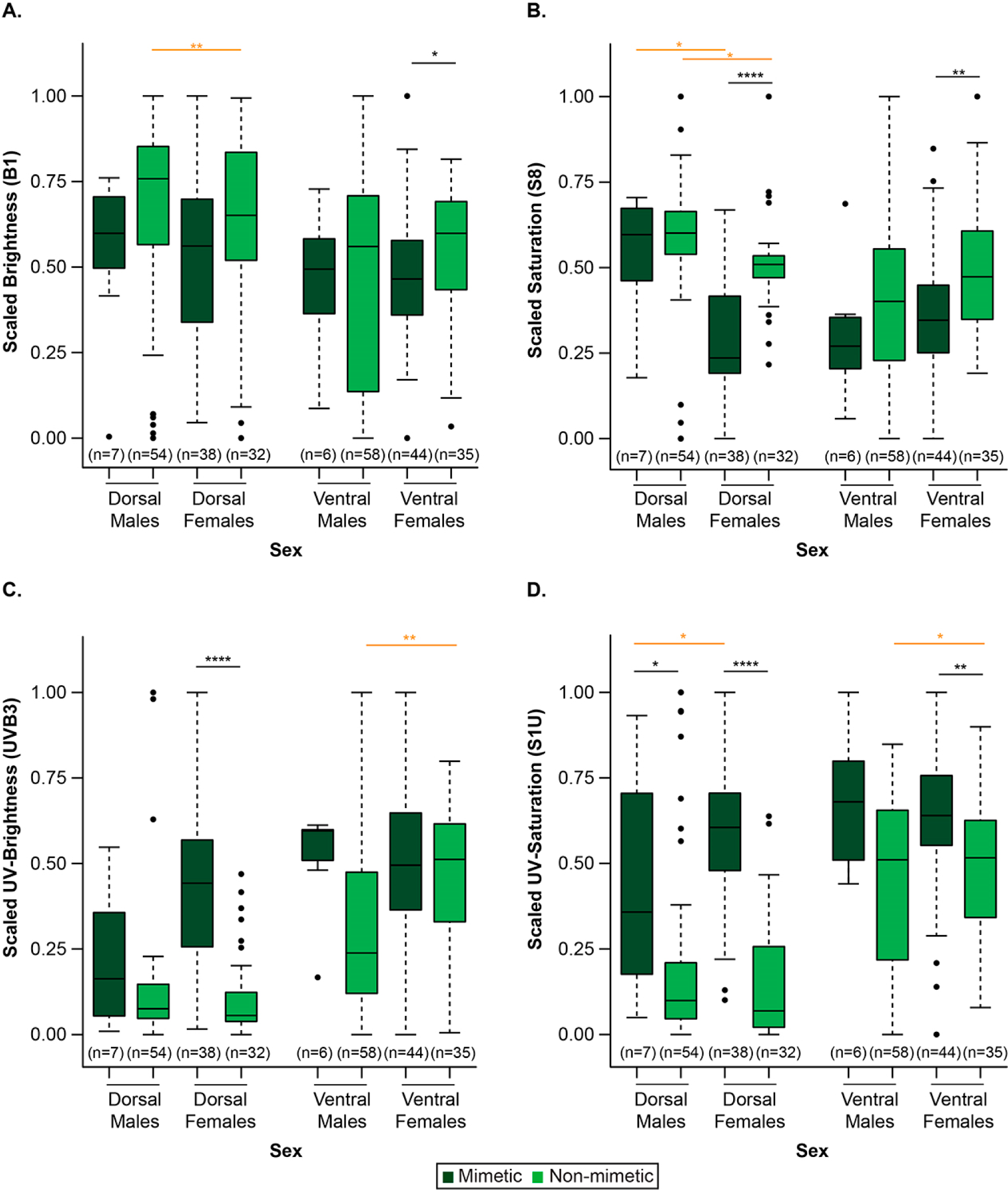
Comparison between white patches of mimetic and non-mimetic butterflies. **A:** brightness, and **B:** saturation, of bright white (mimetic butterflies) and creamy white/yellow/green patches (non-mimetic butterflies) across the entire spectral range (300 nm–700 nm). **C:** brightness, and **D:** saturation, of bright white and creamy white/yellow/green patches across the UV range (300 nm– 400 nm). Black lines and stars represent statistical significance among non-mimetic males and females versus mimetic males and females. Orange lines and stars represent statistical significance among non-mimetic males versus non-mimetic females, and mimetic males versus mimetic females. Brightness values are normalized from 0 to 1 based on min-max readings. Black dots represent outliers. Statistical significance of T-tests or Mann-Whitney tests: *: p<0.05, **: p<0.01, ***: p<0.001, ****: p<0.0001.

These comparisons indicated that creamy white/yellow/green patches of non-mimetic females may be more conspicuous than the bright white patches of mimetic females especially against the brighter black background. Additionally, mimetic males did not differ in brightness or saturation from non-mimetic males, which may suggest a constraint imposed by female choice on male colouration.

### Non-mimetic Males Had Brighter and More Saturated Colour Patches on Dorsal Wing Surfaces Compared with Non-Mimetic Females

Non-mimetic males had brighter and more saturated colour patches on dorsal surfaces (Table S7B, orange lines and stars in Fig 4A, and yellow boxes in Table S9B). Interestingly, UV brightness was higher on the ventral surface in non-mimetic females compared with non-mimetic males (Table S7B, orange lines and stars in Fig 4C, and yellow boxes in Table S9C). This suggests that the UV component on the ventral surface in non-mimetic females might be facilitating the appearance of a more ‘mimetic’ phenotype, as these surfaces also reflect more UV than dorsal surfaces (Fig. 3C). Females had greater UV saturation than males (Table S7B, orange lines and stars in Fig. 4D, and yellow boxes in Table S9C). These differences in the UV range suggest that UV-reflective patches may have a functional role, perhaps in improving mimetic resemblance, that is not particularly well-studied in *Papilio* swallowtails.

### Papiliochrome-II Pigment is the Chemical Basis of Wing Colour Patches in *Papilio*

Prior chemical analyses using Thin Layer Chromatography (TLC), Nuclear Magnetic Resonance (NMR) and preliminary LCMS have suggested that papiliochrome-II pigment was responsible for the creamy white/yellow/green colour patches of *Papilio* (Umebachi 1985; Nishikawa et al. 2013; Yoda et al. 2021). We confirm this with a detailed LCMS analysis in two species with contrasting creamy white (*P. polytes* males) and yellow patches (*P. demoleus*) and bright white mimetic patches (*P. polytes* female form *polytes*). From the extracted ion chromatogram (XIC), we identified a peak at a retention time of ∼18 min that corresponded to the mass of papiliochrome-II (m/z 431.19284) in both *P. polytes* and *P. demoleus* male (Fig.5B, D) samples. We observed this peak in the bright white sample of mimetic females of *P. polytes* as well, but the peak was almost 1000-fold lower, indicating that the papiliochrome-II quantity was negligible (Fig.5C). This pattern was also present in the normalised total ion chromatogram (TIC) and extracted ion chromatogram (XIC) of the three samples for both papiliochrome-II (Fig. 5E: i–iii) and a prominent daughter ion with m/z 152.0707 (Fig. 5E: iv–vi). The *P. demoleus* sample had the highest intensity followed by male *P. polytes* and female *P. polytes f. polytes* (Fig 5B–E). This detailed LCMS analyses confirmed that presence of papiliochrome-II in large quantities was responsible for the creamy white/yellow/green ornamental patches of non-mimetic *Papilio*, whereas highly reduced quantity of papiliochrome-II was responsible for the bright white patches of mimetic butterflies.

**Figure 5:**
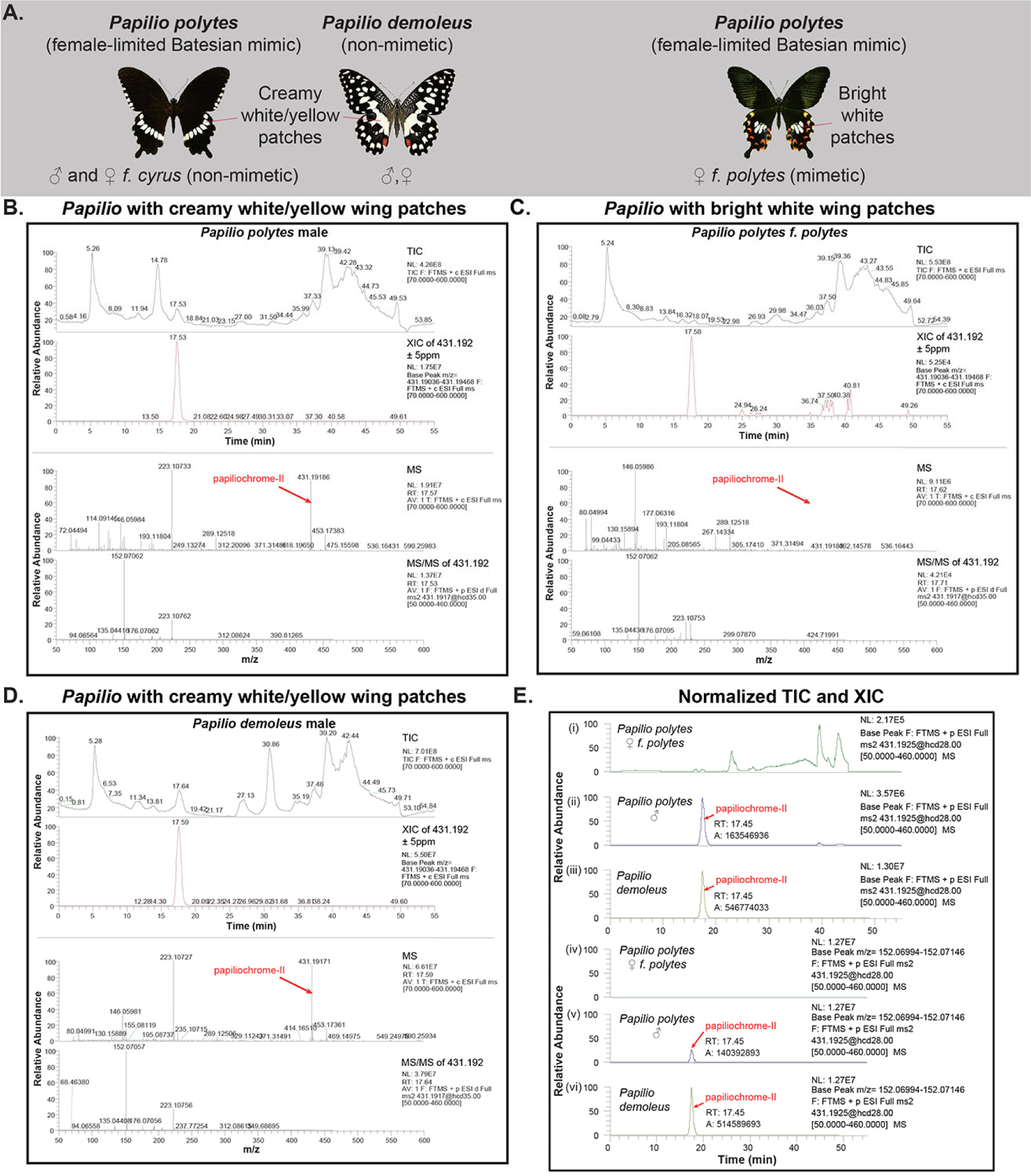
Chemical analysis (LCMS) of wing colour patches of mimetic and non-mimetic butterflies. **A:** Bright white versus creamy white/yellow colour patches of non-mimetic and mimetic butterflies marked on representative images. **B–D:** LCMS peaks of the colour patches of non-mimetic male of *P. polytes* (B), mimetic female *f. polytes* of *P. polytes* (C), and non-mimetic male of *P. demoleus* (D). Red arrows indicate mass peaks for papiliochrome-II pigment. Panels i–iii in **(E)** are stacked total ion chromatograms (TIC) of the three samples with peaks for papiliochrome-II indicated, and iv–vi are the extracted ion chromatograms (XIC) for a major product ion with mass 152.0707, confirming the presence of papiliochrome-II pigment.

## Discussion

In this study we characterised colouration of *Papilio* butterflies with specific interest in the black/brown wing background and colour patches, to understand the nature and extent of variation. This work has not only revealed how spectral properties of colour patches vary across different parameters such as sex, dorsal versus ventral wing surfaces, and mimetic versus non-mimetic forms, but also suggested many interesting hypotheses about functional roles of the colour patches that appear to have shaped the colour pattern variation. Experimental tests of these hypotheses will need to be carried out to confirm that variation in colouration corresponds with their purported functional roles. Such an exercise will be valuable in understanding the evolution of colour signals in this iconic and diverse butterfly genus.

Black/brown background and colour patches varied considerably among the sexes, with males having darker background colouration and brighter colour patches, producing a greater contrast enhancing the colour patches (Fig. 1B–E). This pattern of variation suggests that the colour patches may function as sexual ornaments (Robertson and Monteiro 2005; Kemp 2007), which are more prominent in male than female *Papilio*. However, the fact that both the sexes have these ornamental patches (except in female-limited mimics) suggests that the patches may play a role in species recognition as well as mutual sexual selection (Wallace 1897; Ellers and Boggs 2003; Kunte 2009a). Within this framework, the evolution of brighter and more saturated male patches may have been shaped by stronger female mate choice, whereas relatively less saturated and less bright female patches may have been shaped by natural selection on female patterns to make them less conspicuous in the eyes of predators. Thus, the presence of these colour patches in both the sexes but their more prominent expression in males may suggest a joint action of sexual and natural selection.

We identified brightness as a key spectral parameter that varies between males and females (Figs. 1A–B, 4A, S4A). It is possible that brightness affects mate choice as well as conspicuousness of butterflies to predators, although their relative importance and constraints for the evolution of sexual ornaments and protective colouration remain to be tested. For example, in *Hypolimnas bolina* the bright and white patches on the dorsal surface, which are also highly UV reflective, are used in courtship but may also increase conspicuousness to predators, resulting in males displaying the patches selectively during courtship by manipulating the position of wings (White et al. 2015).

As expected of the courting sex (usually male), our analysis revealed that dorsal wing surfaces of male *Papilio* (*Menelaides*) had higher contrast and saturation compared to ventral wing surfaces, presumably to enhance their sexual ornaments; however, their colour patches were equally bright on both the wing surfaces (Fig. 2A–D, Fig. S2A, B). We also found that non-mimetic females had brighter ventral colour patches (Fig. 3A). A possibility that is often overlooked in studies on butterflies, is that females in sexually monomorphic species are subject to mutual sexual selection where males may also prefer brighter females (Tigreros et al. 2014; Huq et al. 2019). A study in *Bicyclus anynana* identified male preference for bright ventral UV-reflective eyespots of females (Huq et al. 2019), suggesting a sexual signalling role for ventral patches in females. Therefore, studies bridging the gaps between male mate choice and mimicry on spectral variation in females may shed light on evolution of sexual ornaments in females, although they may appear to be less prominent. Although male mate-choice for females has been documented (Chouteau et al. 2017; Westerman et al. 2018), conducting detailed mate-choice experiments with suitable wing manipulations can address how changing brightness, saturation, or contrast of patches on either wing surface may influence mating success and mimetic resemblance in both the sexes, but especially in females.

With several *Papilio* species and female forms exhibiting mimicry, variation in colouration is also structured by natural selection for protective colouration. We characterized variation of black/brown background and white/yellow/green colours of mimetic and non-mimetic forms, which revealed that brightness of colour patches did not vary between wing surfaces except that non-mimetic females had brighter patches on ventral wing surfaces (Fig. 3A). We also observed that UV brightness and saturation were higher on the ventral side (Fig. 3C–D) and mimetic females were more UV reflective on dorsal wing surfaces than non-mimetic females (Fig. 4C). UV reflectance in butterflies has multiple functions, for instance, variations in the dorsal UV region of wing colour spectra are utilized for conspecific identification in *Heliconius* mimicry rings (Dell’Aglio et al. 2018). UV colour signals may also be used for courtship by male (Papke et al. 2007; Rutowski et al. 2007) and female butterflies (Huq et al. 2019). These studies highlight the functional significance of seemingly similar colour patterns that possess relatively subtle variations, which are perceivable by butterflies but not predators. Indeed, cryptic variation in colour spectra and contrast between black and yellow patches in *Heliconius* butterflies (Bybee et al. 2012) and some of their *Melinaea* co-mimics (Llaurens et al. 2014) are used in species recognition in addition to mimetic signals, where patches contribute differentially to the two functions.

It is possible that non-mimetic females have higher UV reflectance on ventral wing surfaces compared with dorsal surfaces (Fig. 3C, Fig. 4C) because the ventral patches function as sexual signals for males (females are often perched with wings closed over their backs while males are courting). Is it possible that this predisposition to be more UV reflective on ventral surfaces may have generated a somewhat pre-adapted phenotype (i.e., more UV-reflective patches on ventral surfaces) that could easily evolve further in mimetic females, which reflect even greater amount of UV on their ventral patches, which perhaps has a role in Batesian mimicry (Fig. S3, 4A). These differences in coloration between the two wing surfaces and mimetic forms may also be attributed to signal partitioning, where distinct functions of mimicry and mate choice are accommodated on different wing surfaces. For instance, ventral wing surfaces exhibit a greater degree of mimetic resemblance among female Batesian mimics compared with their dorsal surfaces as well as compared with males, a pattern that was found across three families of butterflies, including Papilionidae (Su et al. 2015). Even in our present study of the entire clade, male patches of *Papilio* (*Menelaides*) were usually similar on the two wing surfaces (Fig. 2A, 3A). It is possible that although males may benefit from mimicry, the advantage is smaller in males compared with females, and that their sexual ornaments may be under stronger stabilising selection such that their colour patches do not vary significantly across wing surfaces. These sexual differences further bolster the view that non-mimetic males of *Papilio* remain non-mimetic because of a combination of relatively lower mimetic advantage and a stronger selection on sexual ornaments, whereas females are able to evolve towards mimetic phenotypes more readily due to a greater mimetic advantage and weaker stabilising selection on sexual ornaments (Kunte 2008). The greater mimetic advantage to females results from their sexual role—carrying heavy egg-loads and therefore having a constrained escape flight—because of which their mimetic advantage outweighs any potential costs of mimicry; whereas males benefit less from mimicry because they are not as vulnerable to aerial predation owing to their greater flight speeds (Kunte 2009a). Experimentally testing these possibilities may shed light on the functional significance of these consistent colour variations in *Papilio* swallowtails. Such sex-specific differences, female-biased variation, and constrained male phenotypes on different wing surfaces have recently been demonstrated in a number of butterfly species, in the context of mimicry (Su et al. 2015; Basu et al. 2023) as well as outside of it; for example, for mate choice in *Pieris rapae* (Morehouse and Rutowski 2010), thermoregulation in the Himalayan *Pieris canidia* (Gautam and Kunte 2020), and migratory behaviour in milkweed and emigrant butterflies (Bhaumik and Kunte 2018, 2020).

Further, we found that spectral parameters of the black/brown background did not vary between mimics versus non-mimics or mimetic males versus females (black lines and stars in Fig. S4A–B). However, female non-mimics had brighter and more saturated colour patches than mimics when comparing the whole spectral range whereas mimics were brighter and more saturated in the UV region (black lines and stars in Fig. 4A–D). The observed differences in spectral parameters between mimics and non-mimics, as well as the similarities within male and female mimics, likely arise from the selective pressure to enhance mimetic resemblance especially in females. Functional (signalling) consequences of these differences may be tested through visual modelling (Su et al. 2015; Thurman and Seymoure 2016; Dell’Aglio et al. 2018) and predation experiments using mimics, non-mimics, and models (Palmer et al. 2018). Similar effects of variation in brightness and saturation can be tested in *Papilio* butterflies, a system where sexual selection (through courtship and mate choice) and natural selection (through mimetic colouration) potentially affect sex and wing surfaces differently, highlighting the processes that drive wing colour evolution and variation.

Finally, consistent with previous work (Umebachi 1985; Nishikawa et al. 2013; Yoda et al. 2021), we identified papiliochrome-II as the pigment responsible for the creamy white/yellow/green patches that are widely found in non-mimetic butterflies. However, in contrast to these previous studies, we were able to show for the first time with more sensitive LCMS instruments and thorough chemical analysis, that even mimetic butterflies produce papiliochrome-II pigment in the mimetic bright white patches, albeit in trace amounts (Fig. 5). The previous genetic manipulation experiments and their conclusions should be viewed in this light. With a combined strength of the previous genetic manipulations and our detailed chemical analysis, the mechanistic basis of the white/yellow/green sexual colour ornaments of *Papilio* is now much better characterised. This improved understanding may open up further avenues to study the evolution of colouration as sexual ornaments and mimetic signals, and the underlying variation in colouration, in the charismatic *Papilio* swallowtail butterflies.

## Acknowledgments

We thank Shiyu Su for contributing some of the reflectance data; Matthew Lim for advice on photospectrometry measurements; Radhika Venkatesan, Praveen Vemula and Padma Ramakrishnan for advice on pigment characterisation; Blanca Huertas, Geoff Martin and David Lees (Natural History Museum, London) and Ujwala Pawar (Biodiversity Research Collections) for assistance in museum work; and C-CAMP Metabolomics Facility for chemical analysis. Wild-caught samples used for colour spectra measurements were obtained largely from the NCBS campus and private lands, and from wildlife sanctuaries and national parks in India under the following research and collection permits issued by the state forest departments in Arunachal Pradesh (permit no. CWL/G/13(95)/2011-12/Pt-III/2466-70, dated 16/02/ 2015), Karnataka (permit no. PCCF/WL/E2/CR/227/2014-15, dated 2017/05/26), Meghalaya (permit no. FWC/G/173/Pt-II/474-83, dated 27/05/2014), Nagaland (permit no. CWL/GEN/240/522-39, dated 2012/08/14), and West Bengal (permit no. 2115(9)/WL/4 K-1/13/BL41, dated 2013/11/06; and permit no. 1107/42/2 W-705/18, dated 2018/05/07), for which we thank the Principal Chief Conservator of Forest, Chief Wildlife Wardens, Deputy Conservators of Forest, Wildlife Wardens and field officers of those states. This research was supported by an NCBS graduate student fellowship to BD, and Ramanujan Fellowship (Department of Science and Technology, Science and Engineering Research Board, Government of India) and an NCBS Research Grant to KK.

## Author contributions

BD designed research, collected and analysed the data. RV advised on and supervised chemical analysis. KK conceived the project, guided research, provided specimens, instruments and other resources. BD and KK wrote the manuscript. All authors gave final approval for publication and agree to be held accountable for the work performed therein.

## Supplementary figures and tables

**Table S1: Details of Dataset 1 and Dataset 2, and list of comparisons made, for this work.** See Excel sheet, ‘TableS1_DataSubsetsForDifferentComparisons.xlsx’.

**Table S2: Reflectance spectra values for species and specimens used in this work.** See Excel sheet, ‘TableS2_ReflectanceSpectraDatabase.xlsx’.

**Table S3:** Details of statistical analysis (pairwise sign tests) comparing black/brown background and colour patches on the same wing surfaces between sexes. **B1:** Total Brightness; **S8:** Saturation. Significant differences are marked bold (see Figs. 1A, C and Fig. S1 for further details).

**Dataset 1**
| Colour | Wing Surface | Spectral parameter | Mean (female) | Mean (male) | n | statistic | df | p.adj |
| --- | --- | --- | --- | --- | --- | --- | --- | --- |
| **Black/brown background** | **Dorsal** | B1 | 651.3553 | 455.07461 | 39 | 8 | 39 | **0.0003** |
|  |  | S8 | 3.204492 | 2.9707379 | 39 | 16 | 39 | 0.337 |
|  | **Ventral** | B1 | 791.4776 | 535.56058 | 36 | 4 | 36 | **<0.0001** |
|  |  | S8 | 3.0624 | 3.3487633 | 36 | 26 | 36 | **0.011** |
| **White/yellow/green colour patches** | **Dorsal** | B1 | 12571.81 | 15509.036 | 51 | 43 | 51 | **<0.0001** |
|  |  | S8 | 1.462769 | 1.5334973 | 51 | 24 | 51 | 0.78 |
|  | **Ventral** | B1 | 12392.91 | 13052.114 | 58 | 41 | 58 | **0.002** |
|  |  | S8 | 1.146738 | 1.2977126 | 58 | 45 | 58 | **<0.0001** |

**Dataset 2**
| Colour | Wing Surface | Spectral parameter | Mean (female) | Mean (male) | n | statistic | df | p.adj |
| --- | --- | --- | --- | --- | --- | --- | --- | --- |
| **Black/brown background** | **Dorsal** | B1 | 396.89 | 292.80 | 28 | 21 | 28 | **0.013** |
|  |  | S8 | 2.66 | 2.54 | 28 | 13 | 28 | 0.851 |
|  | **Ventral** | B1 | 703.30 | 502.55 | 28 | 23 | 28 | **<0.001** |
|  |  | S8 | 2.31 | 2.33 | 28 | 15 | 28 | 0.851 |
| **White/yellow/green colour patches** | **Dorsal** | B1 | 16299.95 | 18331.86 | 19 | 4 | 19 | **0.019** |
|  |  | S8 | 1.58 | 1.58 | 19 | 11 | 19 | 0.648 |
|  | **Ventral** | B1 | 17118.90 | 18385.91 | 19 | 5 | 19 | 0.064 |
|  |  | S8 | 1.29 | 1.33 | 19 | 8 | 19 | 0.648 |

**Table S4:** Details of statistical analysis (pairwise sign tests) comparing black/brown background and colour patches on dorsal versus ventral wing surfaces. **B1:** Total Brightness; **S8:** Saturation. Significant differences are marked bold (see Figs. 2A, C and Fig. S2 for further details).

**Dataset 1**
| Colour | Sex | Spectral parameter | Mean (dorsal) | Mean (ventral) | Sample size (n) | Statistic | df | p.adj |
| --- | --- | --- | --- | --- | --- | --- | --- | --- |
| **Black/brown background** | Male | B1 | 414.3794 | 551.9489 | 48 | 14 | 48 | **0.006** |
|  |  | S8 | 2.956145 | 3.173625 | 48 | 22 | 48 | 0.665 |
|  | Female | B1 | 656.3148 | 817.8829 | 34 | 11 | 34 | 0.058 |
|  |  | S8 | 3.250374 | 3.029757 | 34 | 25 | 34 | **0.009** |
| **White/yellow/green colour patches** | Male | B1 | 14362.46 | 14077.4 | 51 | 26 | 51 | 1 |
|  |  | S8 | 1.51698 | 1.274299 | 51 | 42 | 51 | **<0.0001** |
|  | Female | B1 | 12149.98 | 13033.08 | 66 | 21 | 66 | **0.004** |
|  |  | S8 | 1.387379 | 1.108886 | 66 | 59 | 66 | **<0.0001** |

**Dataset 2**
| Colour | Sex | Spectral parameter | Mean (dorsal) | Mean (ventral) | Sample size (n) | Statistic | df | p.adj |
| --- | --- | --- | --- | --- | --- | --- | --- | --- |
| **Black/brown background** | Male | B1 | 314.14 | 536.42 | 40 | 5 | 40 | **<0.0001** |
|  |  | S8 | 2.19 | 2.03 | 40 | 24 | 40 | 0.268 |
|  | Female | B1 | 421.13 | 714.02 | 38 | 3 | 38 | **<0.0001** |
|  |  | S8 | 2.66 | 2.28 | 38 | 29 | 38 | **0.002** |
| **White/yellow/green colour patches** | Male | B1 | 17493.12 | 17492.9 | 25 | 14 | 25 | 0.69 |
|  |  | S8 | 1.5 | 1.26 | 25 | 25 | 25 | **<0.0001** |
|  | Female | B1 | 15581.49 | 16539.81 | 24 | 8 | 24 | 0.152 |
|  |  | S8 | 1.48 | 1.23 | 24 | 24 | 24 | **<0.0001** |

**Table S5:** Details of statistical analysis (pairwise sign tests) comparing white colour patches on dorsal versus ventral wing surfaces of mimics and non-mimics. **B1:** Total Brightness; **S8:** Saturation; **S1U:** UV-chroma; **UVB3:** UV-Brightness. Significant differences are marked bold (see Fig. 3A–D for further details).

| Mimetic Status | Sex | Spectral parameter | Mean (dorsal) | Mean (ventral) | Sample size (n) | Statistic | df | p.adj |
| --- | --- | --- | --- | --- | --- | --- | --- | --- |
| **Mimetic** | Male | B1 | 11459.07 | 12422.13 | 6 | 2 | 6 | 0.687 |
|  |  | S8 | 1.48 | 1.20 | 6 | 4 | 6 | 0.687 |
|  |  | S1U | 0.07 | 0.13 | 6 | 0 | 6 | **0.031** |
|  |  | UVB3 | 10.62 | 20.79 | 6 | 1 | 6 | 0.219 |
|  | Female | B1 | 11363.92 | 12110.26916 | 37 | 14 | 37 | 0.188 |
|  |  | S8 | 1.22 | 1.06 | 37 | 31 | 37 | **<0.0001** |
|  |  | S1U | 0.12 | 0.13 | 37 | 7 | 37 | **<0.001** |
|  |  | UVB3 | 18.67 | 21.21 | 37 | 9 | 37 | **0.003** |
| **Non-mimetic** | Male | B1 | 14749.58 | 14298.10 | 45 | 24 | 45 | 0.766 |
|  |  | S8 | 1.52 | 1.28 | 45 | 38 | 45 | **<0.0001** |
|  |  | S1U | 0.04 | 0.09 | 45 | 2 | 45 | **<0.0001** |
|  |  | UVB3 | 7.51 | 16.60 | 45 | 6 | 45 | **<0.0001** |
|  | Female | B1 | 13152.89 | 14210.46 | 29 | 7 | 29 | **0.008** |
|  |  | S8 | 1.61 | 1.17 | 29 | 28 | 29 | **<0.0001** |
|  |  | S1U | 0.04 | 0.11 | 29 | 1 | 29 | **<0.0001** |
|  |  | UVB3 | 6.96 | 20.48 | 29 | 1 | 29 | **<0.0001** |

**Table S6:** Details of statistical analysis (T-tests or Mann-Whitney tests) for comparing black/brown background (A) and white colour patches (B) of mimics versus non-mimics. **B1:** Total Brightness; **S8:** Saturation; **S1U:** UV-chroma; **UVB3:** UV-Brightness. Significant differences are marked bold (see Figs. 4A–D, S4A–B for further details).

| Sex | Comparison | Wing Surface | Statistical test | Mann-Whitney's U | T-Test (t, df) | p-value |
| --- | --- | --- | --- | --- | --- | --- |
| **Male** | **B1** | Dorsal | Mann-Whitney U | 208 |  | 0.413 |
|  |  | Ventral | Mann-Whitney U | 122 |  | 0.5602 |
| **Male** | **S8** | Dorsal | Unpaired t-test |  | t=0.1061, df=63 | 0.9158 |
|  |  | Ventral | Unpaired t-test |  | t=0.2652, df=52 | 0.7919 |
| **Female** | **B1** | Dorsal | Mann-Whitney U | 134 |  | 0.1782 |
|  |  | Ventral | Mann-Whitney U | 84 |  | **0.0093** |
| **Female** | **S8** | Dorsal | Mann-Whitney U | 149 |  | 0.3578 |
|  |  | Ventral | Unpaired t-test |  | t=1.800, df=35 | 0.0805 |

B)
| Sex | Comparison | Wing Surface | Statistical test | Mann-Whitney's U | T-Test (t, df) | p-value |
| --- | --- | --- | --- | --- | --- | --- |
| **Male** | **B1** | Dorsal | Mann-Whitney U | 108 |  | 0.0675 |
|  |  | Ventral | Mann-Whitney U | 163 |  | 0.8109 |
| **Male** | **S8** | Dorsal | Mann-Whitney U | 171 |  | 0.6966 |
|  |  | Ventral | Unpaired t-test |  | t=0.8705, df=62 | 0.3874 |
| **Male** | **UVB3** | Dorsal | Mann-Whitney U | 141 |  | 0.2882 |
|  |  | Ventral | Mann-Whitney U | 100 |  | 0.0903 |
| **Male** | **S1U** | Dorsal | Mann-Whitney U | 101 |  | **0.0458** |
|  |  | Ventral | Mann-Whitney U | 92 |  | 0.0588 |
| **Female** | **B1** | Dorsal | Mann-Whitney U | 461 |  | 0.0836 |
|  |  | Ventral | Mann-Whitney U | 558 |  | **0.0361** |
| **Female** | **S8** | Dorsal | Mann-Whitney U | 176 |  | **<0.0001** |
|  |  | Ventral | Unpaired t-test |  | t=2.921, df=71.84 | **0.0047** |
| **Female** | **UVB3** | Dorsal | Mann-Whitney U | 122 |  | **<0.0001** |
|  |  | Ventral | Unpaired t-test |  | t=1.053, df=77 | 0.2957 |
| **Female** | **S1U** | Dorsal | Mann-Whitney U | 89 |  | **<0.0001** |
|  |  | Ventral | Unpaired t-test |  | t=2.745, df=76.07 | **0.007** |

**Table S7:**
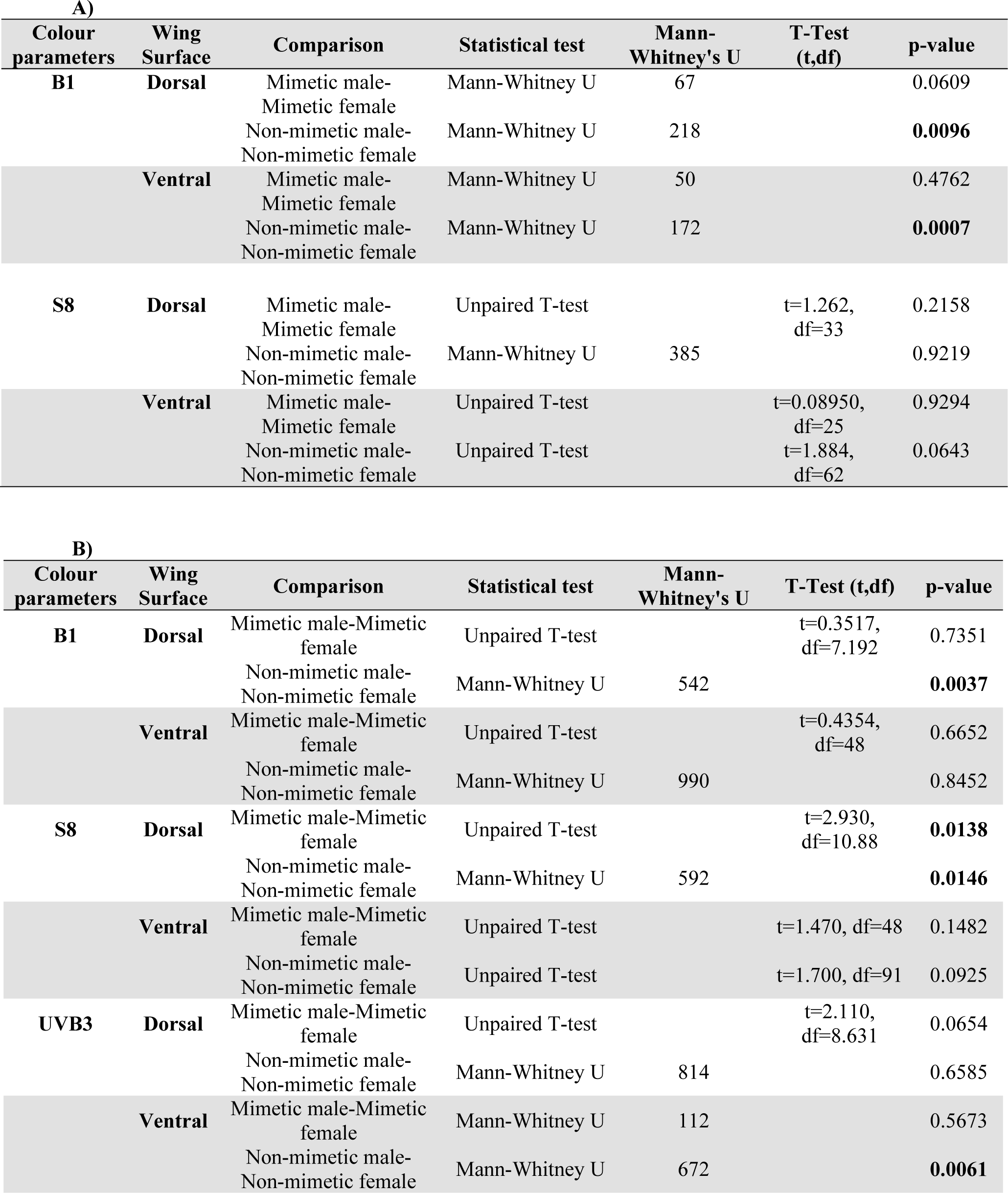
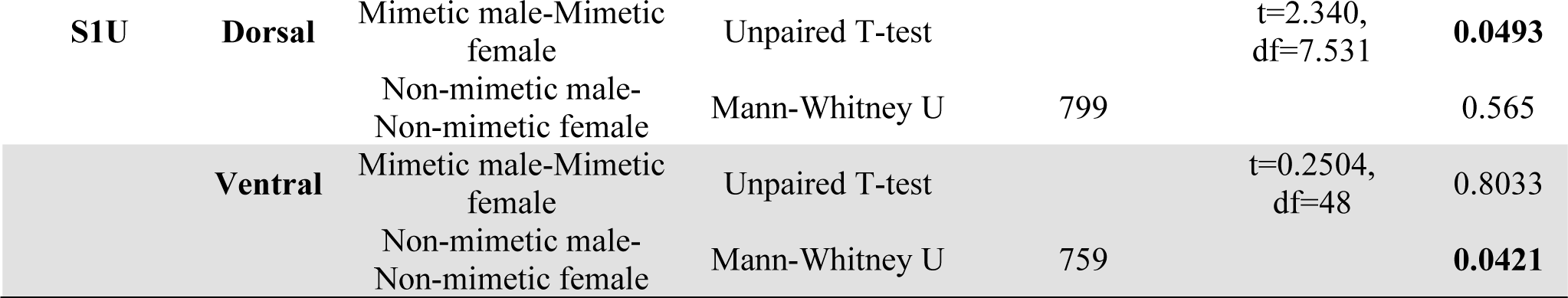
Details of statistical analysis (T-tests or Mann-Whitney tests) for comparing black/brown background (A) and white colour patches (B) of mimetic males versus mimetic females and non-mimetic males versus non-mimetic females. **B1:** Total Brightness; **S8:** Saturation; **S1U:** UV-chroma; **UVB3:** UV-Brightness. Significant differences are marked bold (also see orange lines and stars in Figs. 4A–D and S4A–B).

| Colour parameters | Wing Surface | Comparison | Statistical test | Mann-Whitney's U | T-Test (t,df) | p-value |
| --- | --- | --- | --- | --- | --- | --- |
| **B1** | **Dorsal** | Mimetic male-Mimetic female | Mann-Whitney U | 67 |  | 0.0609 |
|  |  | Non-mimetic male-Non-mimetic female | Mann-Whitney U | 218 |  | **0.0096** |
|  | **Ventral** | Mimetic male-Mimetic female | Mann-Whitney U | 50 |  | 0.4762 |
|  |  | Non-mimetic male-Non-mimetic female | Mann-Whitney U | 172 |  | **0.0007** |
| **S8** | **Dorsal** | Mimetic male-Mimetic female | Unpaired T-test |  | t=1.262, df=33 | 0.2158 |
|  |  | Non-mimetic male-Non-mimetic female | Mann-Whitney U | 385 |  | 0.9219 |
|  | **Ventral** | Mimetic male-Mimetic female | Unpaired T-test |  | t=0.08950, df=25 | 0.9294 |
|  |  | Non-mimetic male-Non-mimetic female | Unpaired T-test |  | t=1.884, df=62 | 0.0643 |

B)
| Colour parameters | Wing Surface | Comparison | Statistical test | Mann-Whitney's U | T-Test (t,df) | p-value |
| --- | --- | --- | --- | --- | --- | --- |
| **B1** | **Dorsal** | Mimetic male-Mimetic female | Unpaired T-test |  | t=0.3517, df=7.192 | 0.7351 |
|  |  | Non-mimetic male-Non-mimetic female | Mann-Whitney U | 542 |  | **0.0037** |
|  | **Ventral** | Mimetic male-Mimetic female | Unpaired T-test |  | t=0.4354, df=48 | 0.6652 |
|  |  | Non-mimetic male-Non-mimetic female | Mann-Whitney U | 990 |  | 0.8452 |
| **S8** | **Dorsal** | Mimetic male-Mimetic female | Unpaired T-test |  | t=2.930, df=10.88 | **0.0138** |
|  |  | Non-mimetic male-Non-mimetic female | Mann-Whitney U | 592 |  | **0.0146** |
|  | **Ventral** | Mimetic male-Mimetic female | Unpaired T-test |  | t=1.470, df=48 | 0.1482 |
|  |  | Non-mimetic male-Non-mimetic female | Unpaired T-test |  | t=1.700, df=91 | 0.0925 |
| **UVB3** | **Dorsal** | Mimetic male-Mimetic female | Unpaired T-test |  | t=2.110, df=8.631 | 0.0654 |
|  |  | Non-mimetic male-Non-mimetic female | Mann-Whitney U | 814 |  | 0.6585 |
|  | **Ventral** | Mimetic male-Mimetic female | Mann-Whitney U | 112 |  | 0.5673 |
|  |  | Non-mimetic male-Non-mimetic female | Mann-Whitney U | 672 |  | **0.0061** |
| **S1U** | **Dorsal** | Mimetic male-Mimetic female | Unpaired T-test |  | t=2.340,<br>df=7.531 | **0.0493** |
|  |  | Non-mimetic male-Non-mimetic female | Mann-Whitney U | 799 |  | 0.565 |
|  | **Ventral** | Mimetic male-Mimetic female | Unpaired T-test |  | t=0.2504,<br>df=48 | 0.8033 |
|  |  | Non-mimetic male-Non-mimetic female | Mann-Whitney U | 759 |  | **0.0421** |

**Table S8:** Summary matrix of results of the pairwise comparisons conducted for this study. **A:** Brightness (B1), and **B:** Saturation (S8). The upper-right of the diagonal represents comparisons for black/brown background (‘Black’) and the lower-left of the diagonal represents comparisons for the colour patches (‘White’). Green boxes are comparisons across the two wing surfaces, yellow boxes are comparisons across the sexes. Acronyms in the box either indicate whether the pairwise differences were not significant (NS) or indicate the category with higher brightness or saturation if the pairwise differences were significant. MD = Male Dorsal, MV = Male Ventral, FD = Female Dorsal, FV = Female Ventral, NA = Comparison Not Applicable. Details of statistical tests are given in Table S3–S4.

**A. Comparison of brightness of black/brown background and colour patches:**
| Brightness (B1)<br>Dataset 1 |  |  |  |  |
| --- | --- | --- | --- | --- |
|  | Male Dorsal | Male Ventral | Female Dorsal | Female Ventral |
| Male Dorsal | White \ Black | MV | FD | NA |
| Male Ventral | NS | White \ Black | NA | FV |
| Female Dorsal | MD | NA | White \ Black | NS |
| Female Ventral | NA | MV | FV | White \ Black |
| Dataset 2 |  |  |  |  |
|  | Male Dorsal | Male Ventral | Female Dorsal | Female Ventral |
| Male Dorsal | White \ Black | MV | FD | NA |
| Male Ventral | NS | White \ Black | NA | FV |
| Female Dorsal | MD | NA | White \ Black | FV |
| Female Ventral | NA | NS | NS | White \ Black |

**B. Comparison of saturation of black/brown background and colour patches:**
| Saturation (S8)<br>Dataset 1 |  |  |  |  |
| --- | --- | --- | --- | --- |
|  | Male Dorsal | Male Ventral | Female Dorsal | Female Ventral |
| Male Dorsal | White \ Black | NS | NS | NA |
| Male Ventral | MD | White \ Black | NA | MV |
| Female Dorsal | NS | NA | White \ Black | FD |
| Female Ventral | NA | MV | FD | White \ Black |
| Dataset 2 |  |  |  |  |
|  | Male Dorsal | Male Ventral | Female Dorsal | Female Ventral |
| Male Dorsal | White \ Black | NS | NS | NA |
| Male Ventral | MD | White \ Black | NA | NS |
| Female Dorsal | NS | NA | White \ Black | FD |
| Female Ventral | NA | NS | FD | White \ Black |

**Table S9:**
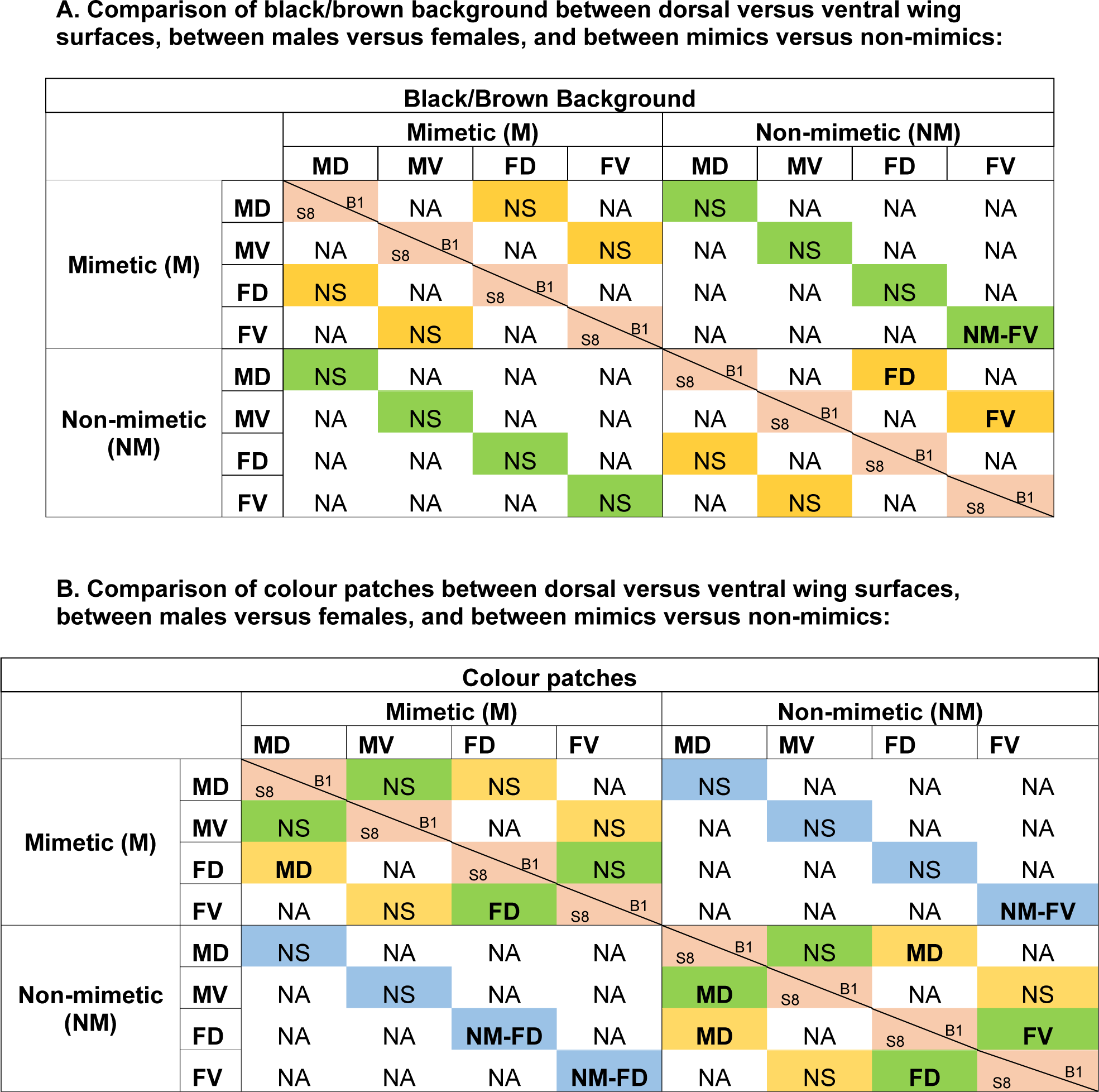
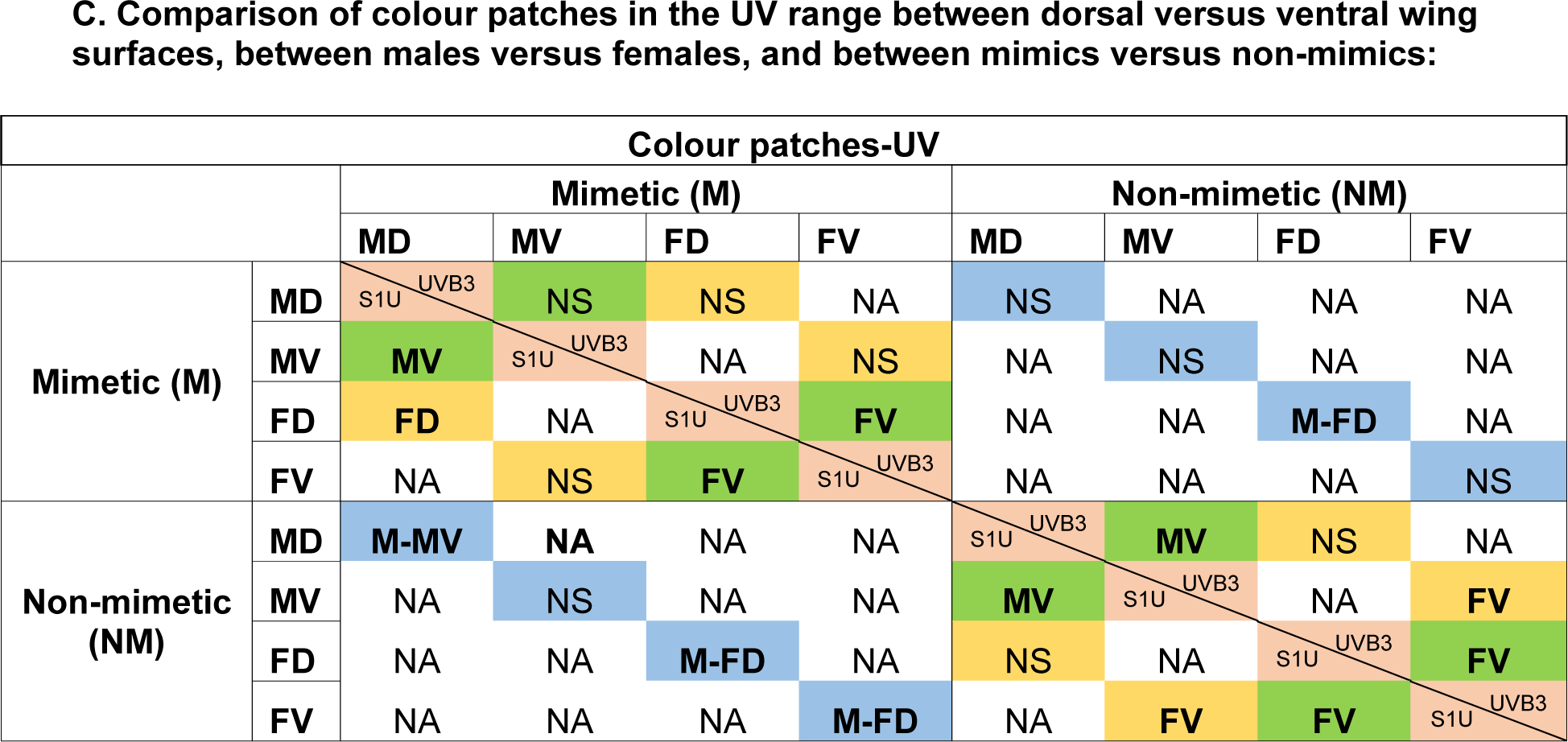
Summary matrix of results of the pairwise comparisons specifically in the context of mimicry. **A:** Black/brown background, **B:** colour patches, and **C:** UV range of colour patches. The upper-right of the diagonal represents comparisons for brightness (B1 or UVB3) and the lower-left of the diagonal represents comparisons for the saturation (S8 or S1U). For **A,** green boxes are comparisons of mimicry versus non-mimicry, yellow boxes are comparisons of males versus females. For **B and C,** green boxes are comparisons across dorsal versus ventral wing surfaces, yellow boxes are comparisons of males versus females, and blue boxes are comparisons of mimicry versus non-mimicry. Acronyms in the box either indicate whether the pairwise differences were not significant (NS) or indicate the category with higher brightness or saturation if the pairwise differences were significant. M = Mimetic, NM = Non-mimetic, MD = Male Dorsal, MV = Male Ventral, FD = Female Dorsal, FV = Female Ventral, NA = Comparison Not Applicable. Details of statistical tests are given in Table S5–S7.

**Figure S1:**
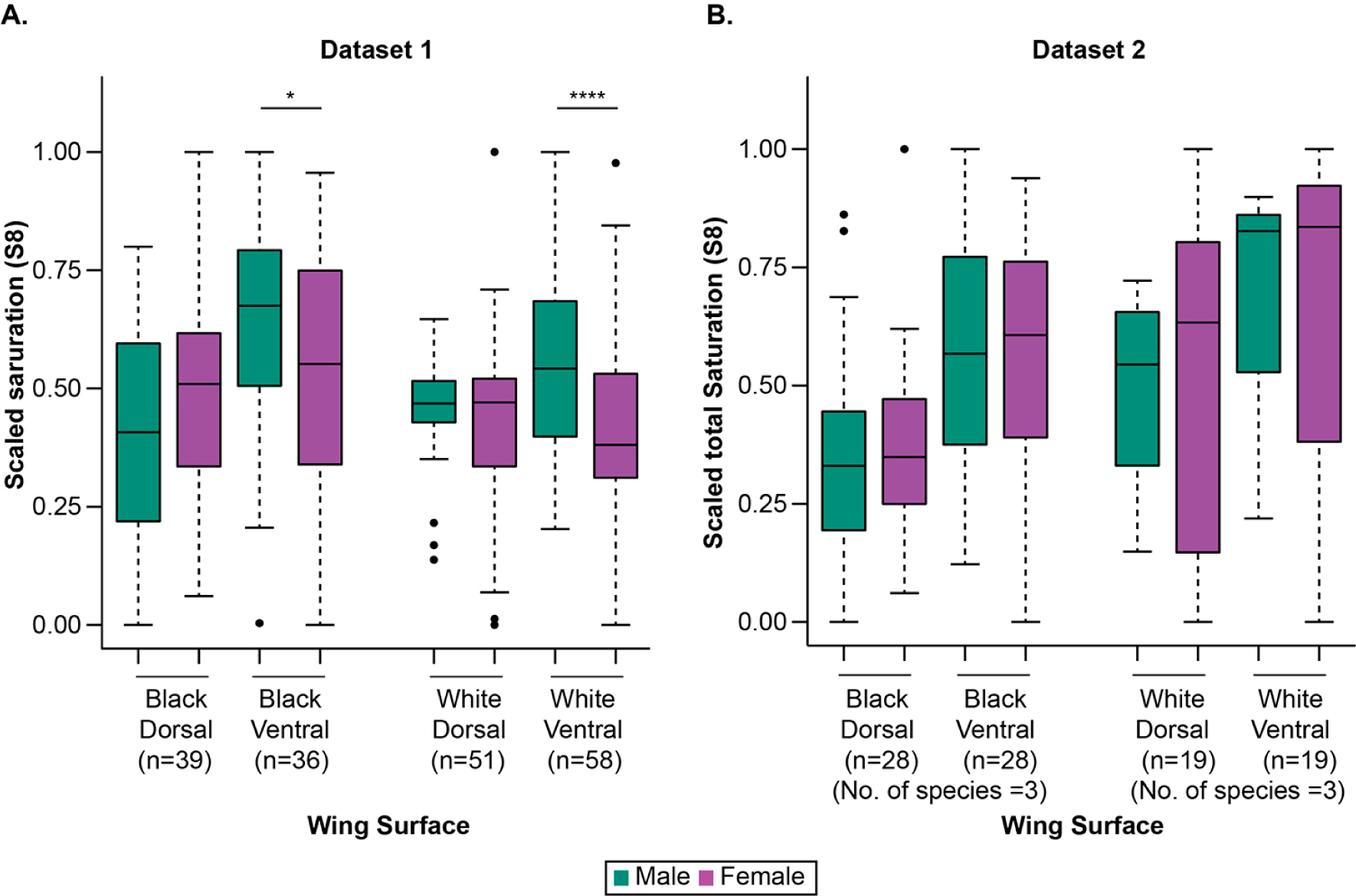
Comparison of black/brown wing background and white patches on the same wing surface between sexes. **A:** Dataset 1; **B:** Dataset 2. Saturation values are normalized from 0 to 1 based on min-max readings. Black dots represent outliers. Statistical significance of pairwise sign tests: *: p<0.05, **: p<0.01, ***: p<0.001, ****: p<0.0001.

**Figure S2:**
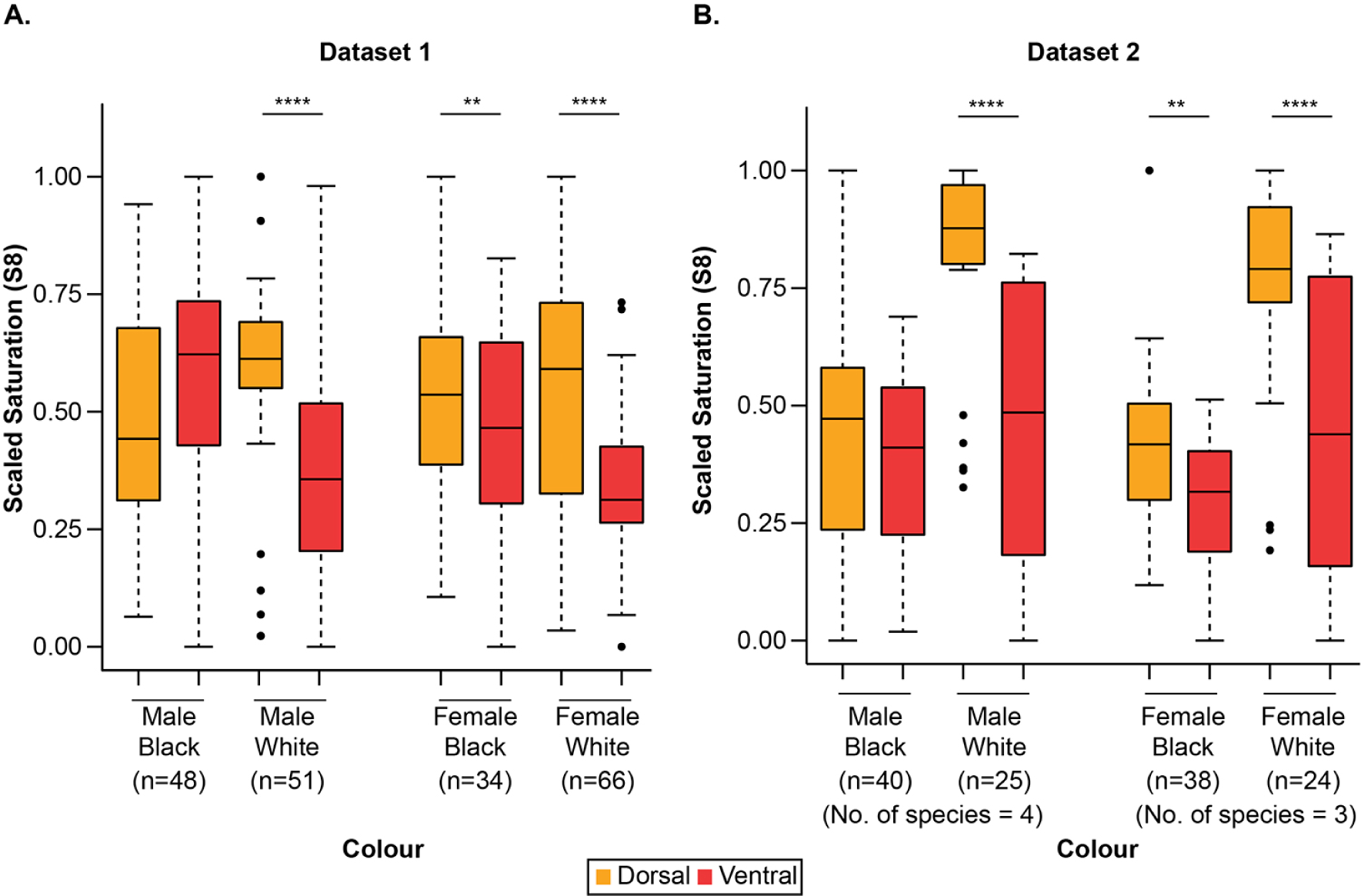
Comparisons of black and white colours within sex across wing surfaces. **A:** Dataset 1; **B:** Dataset 2. Saturation values are normalized from 0 to 1 based on min-max readings. Black dots represent outliers. Statistical significance of pairwise sign tests: *: p<0.05, **: p<0.01, ***: p<0.001, ****: p<0.0001.

**Figure S3:**
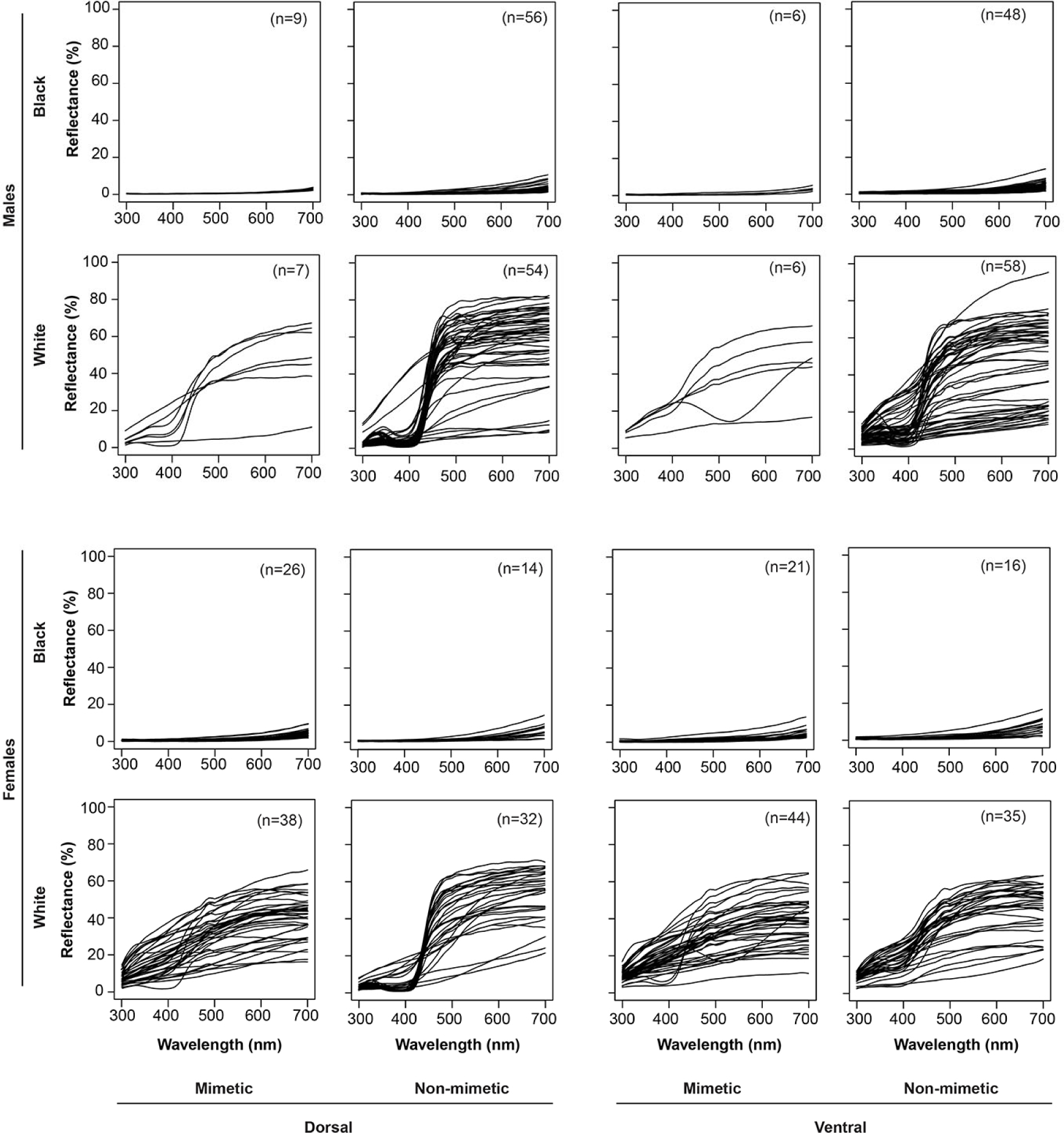
Spectra of black and white wing colours used in colour comparisons in mimetic and non-mimetic butterflies.

**Figure S4:**
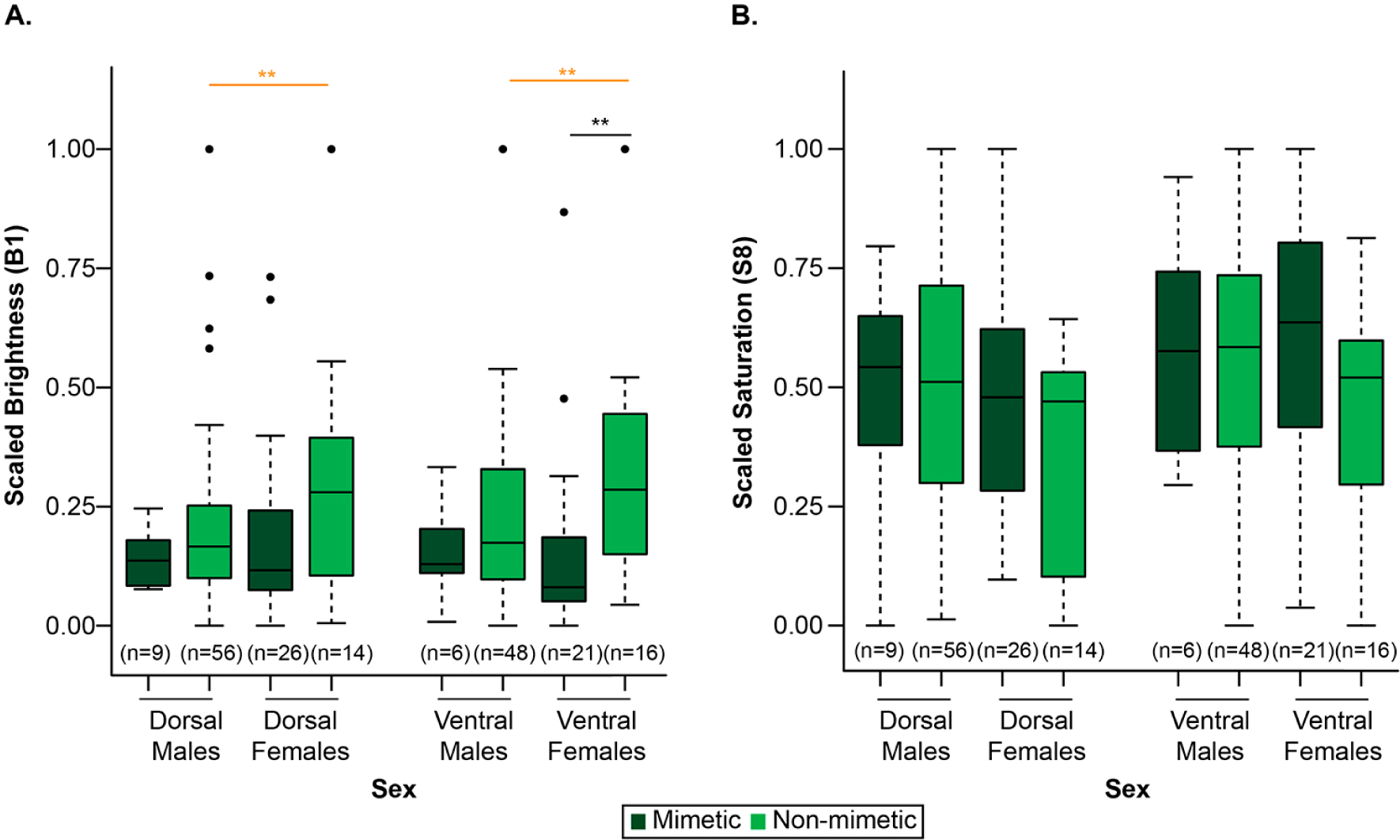
Comparison between mimetic and non-mimetic black patches. **A:** Brightness, and **B:** saturation, of black/brown wing background colour of mimetic and non-mimetic butterflies. Black lines and stars represent comparisons within males or females. Orange lines and stars indicate comparisons between males and females. Brightness and saturation values are normalized from 0 to 1 based on min-max readings. Black dots represent outliers. Statistical significance of T-tests or Mann-Whitney tests: *: p<0.05, **: p<0.01, ***: p<0.001, ****: p<0.0001.

## Notes

### Competing Interest Statement

The authors have declared no competing interest.

